# Sub-tomogram averaging in RELION

**DOI:** 10.1101/030544

**Authors:** Tanmay A.M. Bharat, Sjors H.W. Scheres

**Affiliations:** Structural Studies Division, MRC Laboratory of Molecular Biology, Cambridge CB2 0QH, United Kingdom

## Abstract

Electron cryo-tomography (cryo-ET) and sub-tomogram averaging allow structure determination of macromolecules *in situ,* and are gaining in popularity for initial model generation for single-particle analysis. We describe herein, a protocol for sub-tomogram averaging from cryo-ET data using the RELION software. We describe how to calculate newly developed three-dimensional models for the contrast transfer function and the missing wedge of each sub-tomogram, and how to use these models for regularized-likelihood refinement. This approach has been implemented in the existing workflow for single-particle analysis, so that users may conveniently tap into existing capabilities of the RELION software. As example applications, we present analyses of purified hepatitis B capsid particles and *S. cerevisiae* 80S ribosomes. In both cases, we show that following initial classification, sub-tomogram averaging in RELION allows *de novo* generation of initial models, and provides high-resolution maps where secondary structure elements are resolved.

## Introduction

Imaging macromolecular complexes that are frozen in a thin layer of vitreous ice using an electron microscope (cryo-EM) is rapidly gaining in popularity. A single cryo-EM image may contain 2D projections of many copies of the same complex in different orientations, and these 2D projections may be combined in a 3D reconstruction of its scattering potential. This technique, which is known as single-particle analysis, has recently undergone significant progress with the development of highly efficient direct-electron detectors and improved image processing software. Notably, this technique now allows near-atomic resolution structure to be calculated without the need for crystallisation and from as little as 10-100 μg of purified material (Bai et al., 2015; Cheng, 2015).

Alternatively, in electron tomography (ET) multiple images are taken of the same sample region at different tilt angles in the microscope. From such a series of tilted images, a 3D reconstruction, or tomogram, may be obtained of a unique 3D object such as an entire cell (Baumeister, 2002). Thereby, this technique provides the unique possibility to image macromolecular complexes in their native environment. Moreover, if many copies of a complex of interest are present in tomograms, then the reconstructed 3D density corresponding to each complex may be computationally extracted, and the resulting 3D ‘sub-tomograms’ may be averaged together to increase the signal-to-noise ratio and thereby produce a higher resolution 3D structure (Briggs, 2013). This technique is called sub-tomogram averaging, and it has been successfully applied in numerous cases to reveal biological structures *in situ* or in environments that are otherwise not amenable to single-particle analysis (Beck et al., 2007; Briggs et al., 2009; Förster et al., 2005; Grünewald et al., 2003; Lin et al., 2014; Nicastro et al., 2006).

To date, the use of sub-tomogram averaging is not as widespread as that of single-particle analysis. An important limitation of sub-tomogram averaging is that the best resolved structures by this technique are markedly lower in resolution than those from single-particle analysis (Briggs, 2013). Another reason may be that tomographic data collection is slower, and sub-tomogram averaging requires more complicated image processing, since tomographic reconstruction needs to be followed by alignment and classification of the sub-tomograms. Still, the advantage of being able to study macromolecules *in situ* remains extremely attractive. This is powerfully illustrated by the recent application of sub-nanometer resolution cryo-ET sub-tomogram averaging to the HIV-1 capsid (Schur et al., 2015) and to membrane-bound ribosomes (Pfeffer et al., 2015). Further developments of both experimental data acquisition procedures (Rigort and Plitzko, 2015) and image processing algorithms (Hrabe et al., 2012) will continue to drive this technique towards higher resolutions and wider applicability.

Recently, we introduced a new image processing approach to sub-tomogram averaging (Bharat et al., 2015) that is based on regularized likelihood optimization in the RELION program (Scheres, 2012a, b). This program was originally designed for single-particle analysis and has been used to calculate numerous near-atomic resolution structures (Bai et al., 2015). Because the sub-tomogram averaging approach in RELION was modelled on the single-particle workflow, existing RELION users will find many similarities (Figure 1). The main deviation from the single-particle workflow lies in the generation of a 3D model for the information transfer in each sub-tomogram, which is used to compensate for both the missing wedge as well as the effects of the contrast transfer function (CTF) in the tilt series images (Bharat et al., 2015). A significant effort was made to build on existing tools inside RELION, rather than writing new tools specifically for sub-tomogram averaging. This facilitates transitioning between sub-tomogram averaging and single-particle analysis, and thus naturally supports a hybrid approach (Bartesaghi et al., 2012; Bharat et al., 2014; Bharat et al., 2012).

**Figure 1.**
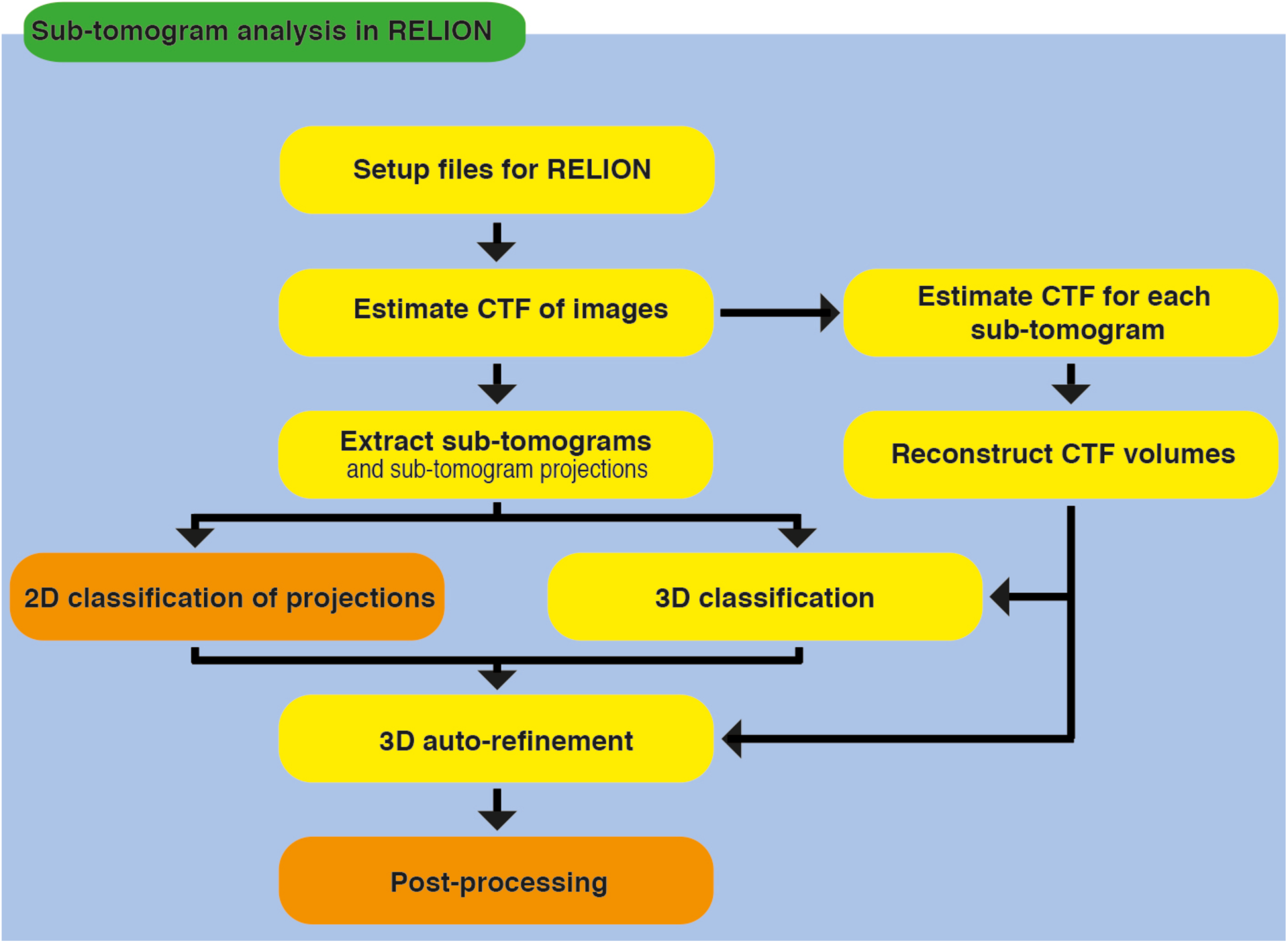
Workflow of the image processing protocol. A schematic representation of the recommended workflow for sub-tomogram analysis using RELION presented in this paper. The main difference between single-particle analysis and subtomogram analysis in RELION is related to CTF estimation and the new 3D CTF model. Steps highlighted in orange are unchanged from the single-particle analysis workflow.

In this protocol we describe the practical use of RELION for sub-tomogram averaging. Our approach complements various single-particle analysis software packages that also offer functionalities for sub-tomogram averaging, e.g. (Heymann and Belnap, 2007; Tang et al., 2007), as well as multiple specialized packages for sub-tomogram averaging, e.g. (Castano-Diez et al., 2012; Förster et al., 2005; Hrabe et al., 2012; Huiskonen et al., 2010; Kremer et al., 1996; Nicastro et al., 2006). As many structure determination projects in practice resort to a combination of different software packages, we will explicitly indicate those points in the workflow that are likely points of conversion between alternative approaches. Recommended procedures for single-particle analysis in RELION are described in detail in the online documentation (the RELION wiki: http://www2.mrc-lmb.cam.ac.uk/relion). Using the protocol described here together with the online documentation, novice users should be able to conduct sub-tomogram averaging for their own project using RELION. We assume basic familiarity with Unix/Linux based systems, and the ability to run provided scripts from the command-line.

## Materials

### Equipment setup

- A computer or a computing cluster with at least 64-128 processors, 120 Gb of shared memory, and 1 Tb of disk space.
- A RELION installation (version 1.4 or newer; available for free from http://www2.mrc-lmb.cam.ac.uk/relion).
- A Python installation (version 2.6 or newer; available from https://www.python.org)
- An IMOD installation (version 4.7 or newer; available for free from http://bio3d.colorado.edu/imod/)

## Procedure

Steps 1-6 explain how input files and directories should be arranged for RELION sub-tomogram averaging.

1| The directory from which all RELION commands and the graphical-user-interface (GUI) is launched will be known as the project directory ( `./`). We recommend making a folder inside the project directory that is called `./Tomograms/` where all the input data is stored.

2| To enter the RELION workflow, we assume that tomograms have been generated in MRC format (Crowther et al., 1996). Tomogram calculation is not done inside RELION, but relies on software packages like IMOD (Kremer et al., 1996), Tomo3D (Agulleiro and Fernandez, 2011), pyTOM (Hrabe et al., 2012) or Bsoft (Heymann and Belnap, 2007). In the examples below, we used the IMOD package for tilt series alignment, and we used Tomo3D for tomographic reconstruction. Generate a sub-directory in the `./Tomograms/` directory with the name of each tomogram in the data set, and copy all tomograms to their respective sub-directories. For example, if there are two tomograms in the data set, then the locations of those tomograms should be as follows -

~~~
./Tomograms/tomogram1/tomogram1.mrc
./Tomograms/tomogram2/tomogram2.mrc
~~~

3| The coordinates of the centres of all macromolecular complexes, or particles, in a tomogram should be saved in the same sub-directories as the corresponding `.mrc` file. The coordinate files should have the suffix `.coords` and the prefix should be the same as the prefix of the tomogram name. In the results presented here we either used the MolMatch software for template matching (Förster et al., 2005) or manually picked particles from IMOD (Kremer et al., 1996). In our two-tomogram example the name and location of the coordinate files should be -

~~~
./Tomograms/tomogram1/tomogram1.coords
./Tomograms/tomogram2/tomogram2.coords
~~~

The coordinates within each file should be written out in a three-column ASCII format, corresponding to the X,Y,Z position of the centre of each sub-tomogram in pixels. Each coordinate file should contain as many lines as there are particles in the tomogram. For example

~~~
100.0       355.0        200.0
2034.0      1100.0       561.0
3011.0      2539.0       321.0
~~~

4| For CTF estimation, the aligned tilt series should also be placed in the same sub directories. For each tomogram, this should be a single MRC stack, and the suffix of the files should be .`mrcs`. It is important that this is the exact same aligned tilt series as the one used for tomographic reconstruction. Again, in our example these should be called -

~~~
./Tomograms/tomogram1/tomogram1.mrcs
./Tomograms/tomogram2/tomogram2.mrcs
~~~

5| Along with the aligned stack, also provide the final tilt angles from IMOD in a separate text file. Each line in these text files should contain one number corresponding to the final tilt angle assigned to that image during alignment. The order of the lines should correspond to the order of the images in the aligned stack.

~~~
59.44
56.44
53.43
~~~

Copy these angles files into the same directory.

~~~
./Tomograms/tomogram1/tomogram1.tlt
./Tomograms/tomogram2/tomogram2.tlt
~~~

If `.tlt` files with angle values are not provided, then the tilt angles will be read from the extended header of the. `mrcs` file, if this exists, in step 9 (see below).

6| For each aligned tilt series, a text file should be created that lists the tilt angles and the accumulated radiation for each image in the tilt series. This information will be used to calculate a dose-dependent 3D CTF model (Figure 2A-B) that also accounts for radiation-induced damage of the specimen. This text file should have as many lines as there are images in the tilt series. The text file should have two columns: the first one for the refined tilt angle (after tilt-series alignment), and the second one for the total accumulated dose in e−/Å^2^ prior to collecting that image. If you provide the nominal tilt angles rather than the refined tilt angles, the python setup script in step 9 will assign accumulated dose values to the closest refined tilt angle from the `.tlt` file. For example –

**Figure 2.**
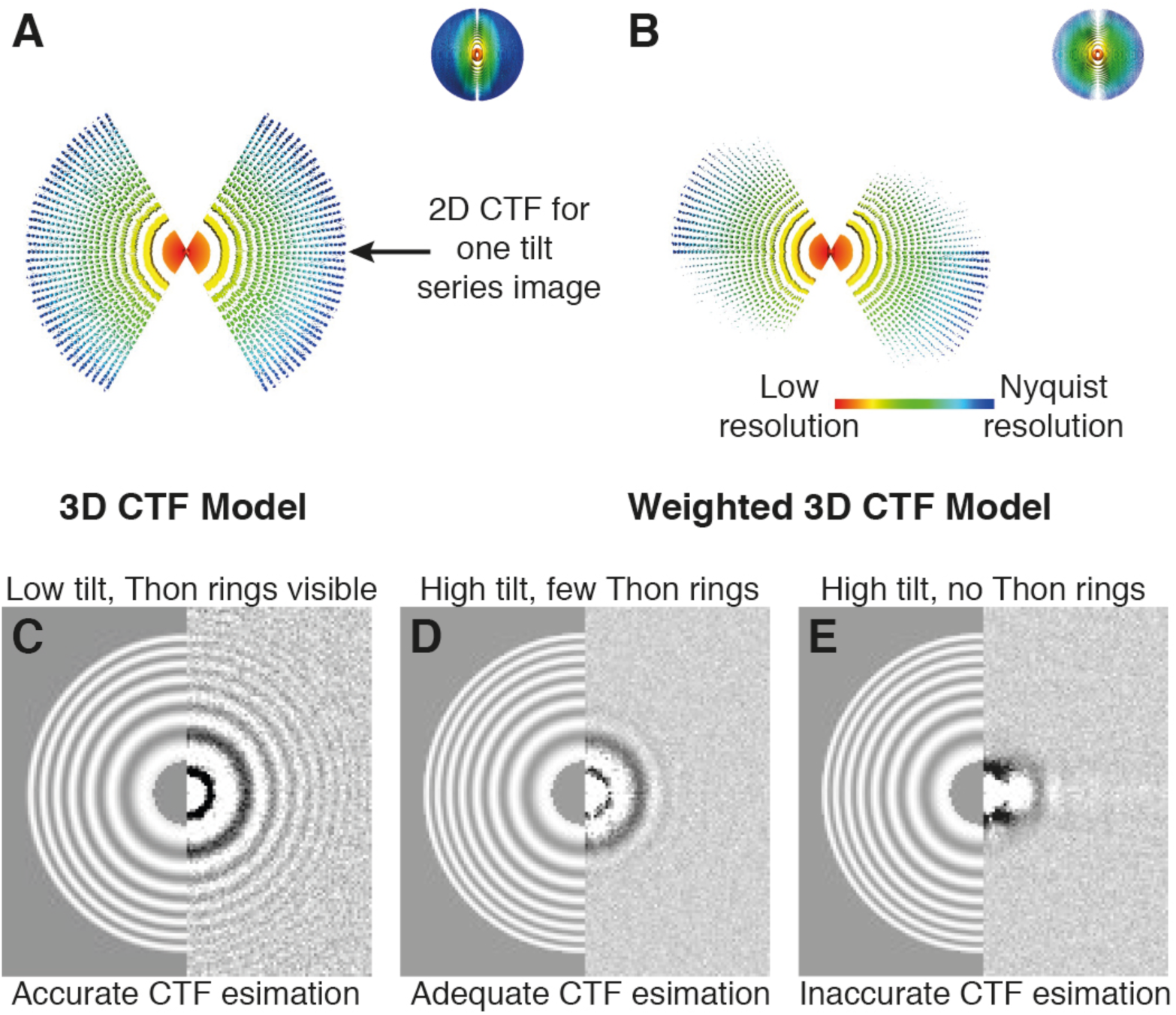
CTF estimation for the 3D CTF model. (A) The unweighted 3D CTF model used in RELION. This model also compensates for the missing wedge. (B) The weighted 3D CTF model used in RELION. The weighted model accounts for increase in noise at high tilts, and for radiation-induced damage. (C) A diagnostic output file of CTFFIND3 from a low-tilt tilt series image. There is no visible radiation-induced motion, and many Thon rings are visible making CTF estimation accurate. (D) Corresponding diagnostic file from a high-tilt image. Less Thon rings are visible die to increased specimen thickness. CTF estimation in this case is adequate but not as accurate as C. (E) A diagnostic file from a high-tilt image where no Thon rings are visible. CTF estimation is not possible from this image, and it should either be removed from the tilt series or the data collection strategy should be modified to include the recording of additional images for CTF estimation on either side of the target region (Bharat et al., 2015).

~~~
61.1       0.0
57.3       2.0
54.2       4.0
~~~

Save these files in the same sub-directories with the suffix `.order` -

~~~
./Tomograms/tomogram1/tomogram1.order
./Tomograms/tomogram2/tomogram2.order
~~~

7| Generate a RELION-type metadata file in the STAR format (Hall, 1991; Scheres, 2012b), called `all_tomograms.star`, using the command line -

~~~
relion_star_loopheader rlnMicrographName > all_tomograms.star
ls./Tomograms/*/*.mrc >> all_tomograms.star
~~~

The output file lists all the tomograms in the data set, and should have the format -

~~~
data_
loop_
_rlnMicrographName
./Tomograms/tomogram1/tomogram1.mrc
./Tomograms/tomogram1/tomogram2.mrc
~~~

Steps 8-14 illustrate how to calculate the 3D CTF models for each sub-tomogram. To facilitate routine application of these steps, we provide a single python script called `relion_prepare_subtomograms.py` on the RELION wiki. This script takes the `all_tomograms.star` file as input, and carries out steps 8-14, depending on the options that the user specifies in its header. For specialized applications or in case of troubleshooting, any of the steps 8-14 may be carried out independently.

8| CTF correction depends on the estimation of a defocus value for every image of the tilt series (i.e. for all the individual images in the `.mrcs` stacks mentioned above). We do this by calling CTFFIND (Mindell and Grigorieff) through the wrapper provided in RELION: `relion_run_ctffind`. Executing this program will be done automatically by the `relion_prepare_subtomograms.py` script. Input parameters for CTFFIND should be changed in the header of the python script. Specify inputs like voltage, spherical aberration and detector pixel size based on your data collection experiment.

? TROUBLESHOOTING

9| Next, run the provided python script from the RELION project directory –

~~~
python relion_prepare_subtomograms.py
~~~

As in single-particle analysis, this script will create a `./Particles/Tomograms/` directory with one sub-directory per tomogram. Following CTF estimation using CTFFIND, in each tomogram sub-directory `.star` files corresponding to the CTF model for each sub-tomogram will be written out. A `particles_subtomo.star` file that lists all the sub-tomograms as well as their corresponding 3D CTF models will also be written to the project directory. This file will be the input for 3D classification and 3D auto-refinement in RELION.

? TROUBLESHOOTING

10| It is important to inspect the diagnostic output of CTFFIND for every image of every tilt series in the data set (Figure 2C-E). This may be conveniently done using the ‘Display’ button in the RELION GUI (Figure 3), and selecting the output `_ctffind.star` file for each tomogram, e.g. `./Tomograms/tomogram1/ctffind/tomogram1_ctffind.star`. If CTF estimation failed for some images (see Figure 2E), then `relion_run_ctffind` should be executed again for that image with different parameters. We recommend that either the entire python script should be run again (step 9), or users can look in the `relion_subtomo_commands.txt` file for the relevant commands for individual tilt series and run them again. Alternatively, one could use a CTF estimation program from an alternative program, possibly one that allows manual steering like EMAN2’s e2ctf.py (Tang et al., 2007), to find a defocus that fits the observed power spectrum of the image. In that case, the resulting defocus value should be inserted manually into the output `_ctffind.star` file by changing the corresponding values using a text file editor.

**Figure 3.**
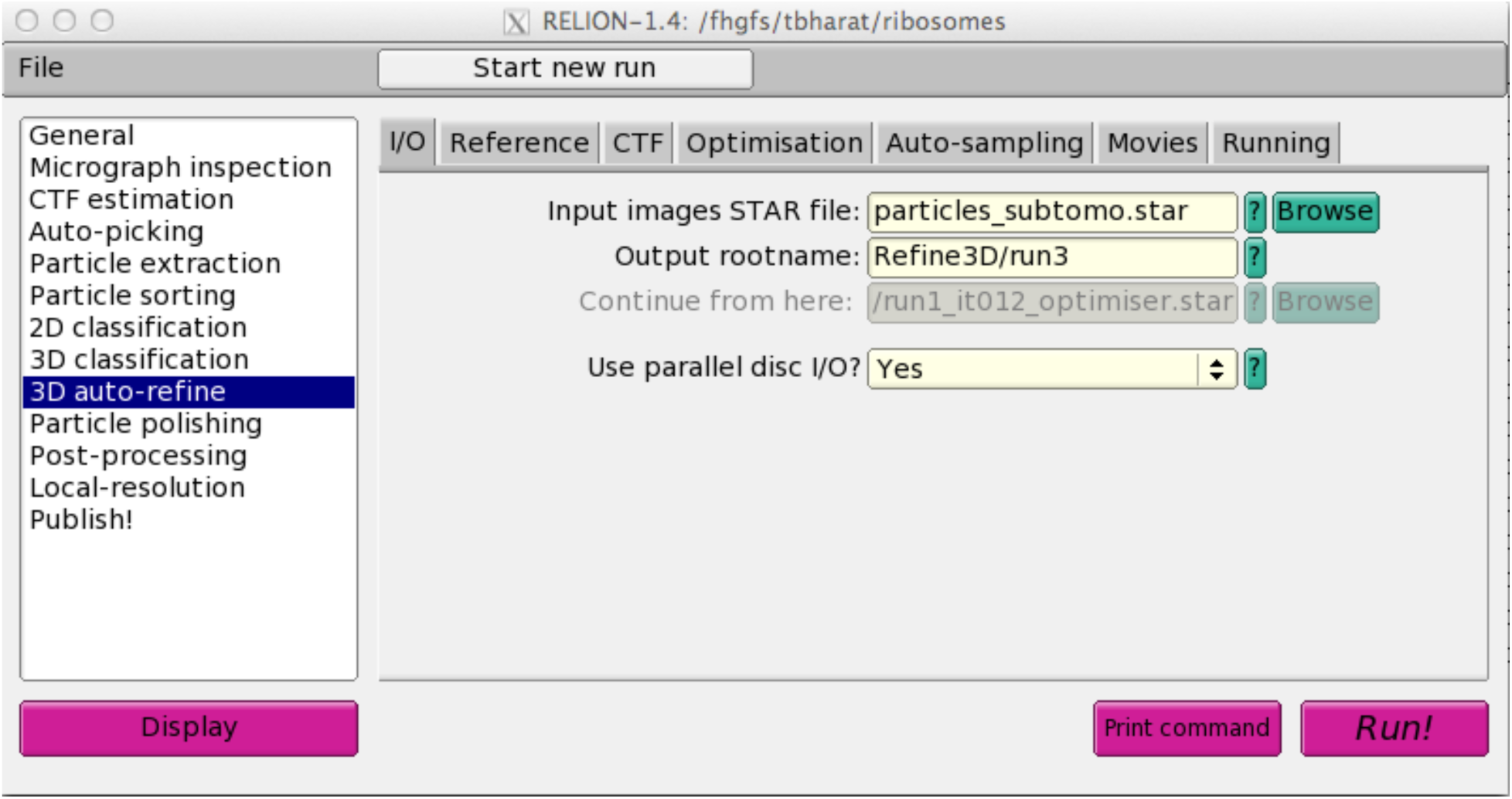
The RELION graphical user interface. After CTF parameters have been estimated for each particle in each image of the tilt series and 3D CTF models have been reconstructed, the actual tasks of sub-tomogram analysis may all be performed using the RELION graphical user interface. The 3D auto-refine page of this user interface is shown. The white column on the left shows different ‘job-types’, which are ordered according to the natural workflow from top to bottom. On the main panel, the ‘**3D auto-refine**’ job-type is shown. This job-type has tabs for “**I/O**”, “**Reference**”, “**CTF**”, “**Optimization**”, “**Auto-sampling**”, “**Movies**”, and “**Running**” where users should enter the input parameters as described in the main text. The “**Display**”, “**Print command**” and “**Run!**” buttons are used to view images, commands and launch jobs, respectively.

? TROUBLESHOOTING

11| *(Steps 11-13 relate to extra options provided to improve CTF estimation accuracy)* For flat samples, the effective thickness of the ice that the electron beam passes through increases with increasing tilt angle. This results in lower signal-to-noise ratios for the higher tilt images, which may preclude reliable CTF estimation. If this is the case, the average defocus measured for the lower tilt images may be applied to the higher tilt images, especially if the applied defocus is stable throughout the tilt series (Schur et al., 2015). This may be done conveniently by setting the `UseOnlyLowerTiltDefoci` variable to `True` in the header of the `relion_prepare_subtomograms.py` script and providing a threshold value for the lower tilt.

12| Another way to estimate the CTF parameters of higher tilt images more accurately is to acquire two extra images for each setting of the stage in the tilt series. We collected these images along the direction of the tilt axis, and spaced equally on either side (Bharat et al.; Eibauer et al., 2012) of the region of interest. If such images were collected, then they should be saved in MRC format with a `.trial` suffix in the relevant tomogram sub-directories -

~~~
./Tomograms/tomogram1/tomogram1.trial
./Tomograms/tomogram2/tomogram2.trial
~~~

Note that the number of images in the `.trial` stack should be exactly double the number of images in the original tilt-series stack and the order of the images should be the same as the aligned tilt series (60°,60°,57°,57° and so on). We do not recommend collecting a single extra image one side of the region of interest because there could be a systematic focus offset between the extra image and the tilt series image. In order to run CTFFIND on these extra images, one should set the `UseTrialsForCtffind` variable in the `relion_prepare_subtomogram.py` script to `True`.

13| The last method to improve the 3D CTF model is to apply a linear, dose-dependent B-factor to the data (also see (Bharat et al., 2015)). Based on observations made for single-particle data sets (Scheres, 2014), we increased the B-factor by 4 Å^2^ for each 1 e^−^/Å^2^ accumulated dose for both examples described in this paper. Users may want to select a different value depending on the radiation sensitivity of their specimen. To set this parameter, change the `Bfactor` variable in the header of the python script. Note that the tilt-dependent scale factor and the position-dependent defocus of each particle will be calculated automatically for every image in the tilt series.

14| The command to reconstruct each 3D CTF model is written out in a run script called `do_all_reconstruct_ctfs.sh`, which is automatically generated in step 9. This script should be run from the RELION project directory –

~~~
./do_all_reconstruct_ctfs.sh 200
~~~

In the above command, the parameter ‘`200`’ determines the size of the generated CTF volumes in pixels. This should be set to the same values as the “Particle box size” in step 15, i.e. the size of the extracted sub-tomograms. The above script contains multiple commands, and may thus be split into shorter text files for convenient parallelization.

15| Next, one extracts every particle into a sub-tomogram by performing a 3D window operation on the corresponding tomogram. On the “**General**” job-type of the RELION GUI (Figure 3), provide the tomogram pixel size in Å, and the diameter of a spherical mask (in Å) that will be used for calculating the average and standard deviation of the background pixel values. Make sure this mask is slightly larger than the longest diameter of the particle. Then, on the “**Particle extraction**” job-type provide the `all_tomograms.star` file (from step 7) as “micrograph STAR file”; set the “Coordinate-file suffix” to `.coords`; and “rootname” should be set to the same as the `RootName` variable in the `relion_prepare_subtomograms.py` script. The “Particle box size” (which is given in pixels) should be set to reflect 150-200% of the particle’s longest dimension. It is recommended to invert the contrast of the sub-tomograms if the particles in the tomogram are black. The re-scaling option may be used to downscale the extracted sub-tomograms, which will save computational resources if the tomograms were taken with a smaller pixel size than necessary for the target resolution. Running this job (by pressing the “**Run!**” button on the GUI) will write out the sub-tomograms as individual `.mrc` files in the `./Particles/Tomograms/tomogram?/` directories that were created by the python setup script in step 9.

16| Along with ordinary sub-tomograms (which are 3D volumes), 2D projections of all sub-tomograms (along the Z-axis) may also be calculated by again using the ‘Particle extraction’ job-type. This is done by running the same job as in step 15, but with providing the extra option --`project3d` on the ‘**Additional arguments**’ line of the ‘**Running**’ tab. By setting the ‘Extract rootname’ on the ‘I/O’ tab to `subtomo_proj2d`, an additional `.star` file called `./subtomo_proj2d.star` will be generated, which lists the 2D projections of all sub-tomograms.

17| The 2D projections may be used in reference-free 2D classification of the data, much like one would use in single-particle analysis, which is a computationally cheaper alternative to classification of the sub-tomograms (Bharat et al., 2015; Yu et al., 2013). On the ‘**I/O**’ tab of the ‘**2D classification**’ job-type, provide the `./subtomo_proj2d.star` as the input `.star` file, and set the ‘Output rootname’ for example to `Class2D/run1`. The number of classes will depend on the number of sub-tomograms and the expected heterogeneity in the data. As a rule of thumb, we typically use at least on average 30 sub-tomograms per class, and we hardly ever use more than 50-100 classes. Because the 2D projections do not have a CTF model, switch CTF-correction off on the ‘**CTF**’ tab. On the ‘**Optimisation**’ tab, we typically perform 25 iterations, and use a regularisation parameter of 2 for 2D classification (Scheres, 2012a). Typically, we mask particles with zeros, and provide a limit on the resolution of around 10-15 Å to include in the expectation (E-) step of the algorithm. On the ‘**Sampling**’ tab, the default parameters are suitable for most cases. Executing this job through the ‘**Run!**’ button will result in multiple output files for each iteration in the `./Class2D` directory. The final class averages are stored in a `.mrcs` stack file called `./Class2D/run1_it025_classes.mrcs`. The `./Class2D/run1_it025_model.star` file contains information about the final classes, such as their relative size and the estimated resolution of each class average.

18| The resulting 2D class averages from step 17 may be visualized using the ‘**Display**’ button on the GUI and selecting the `Class2D/run1_it025_model.star` file. In the subsequent pop-up window, it is useful to reverse-sort the classes on `rlnClassDistribution`, which will place the largest class averages on the top of the display window. At this point, the user needs to select good-looking class averages (which reveal recognizable protein features, e.g. see Figure 4A-B) by double-clicking the class averages, which will put a red border around the image. A new `.star` file with only the particles that correspond to the good classes may then be saved by right-clicking the display window, and selecting the option ‘Save STAR with particles from the selected classes’ option from the pop-up menu as `subtomo_proj2d_sel. star`.

**Figure 4.**
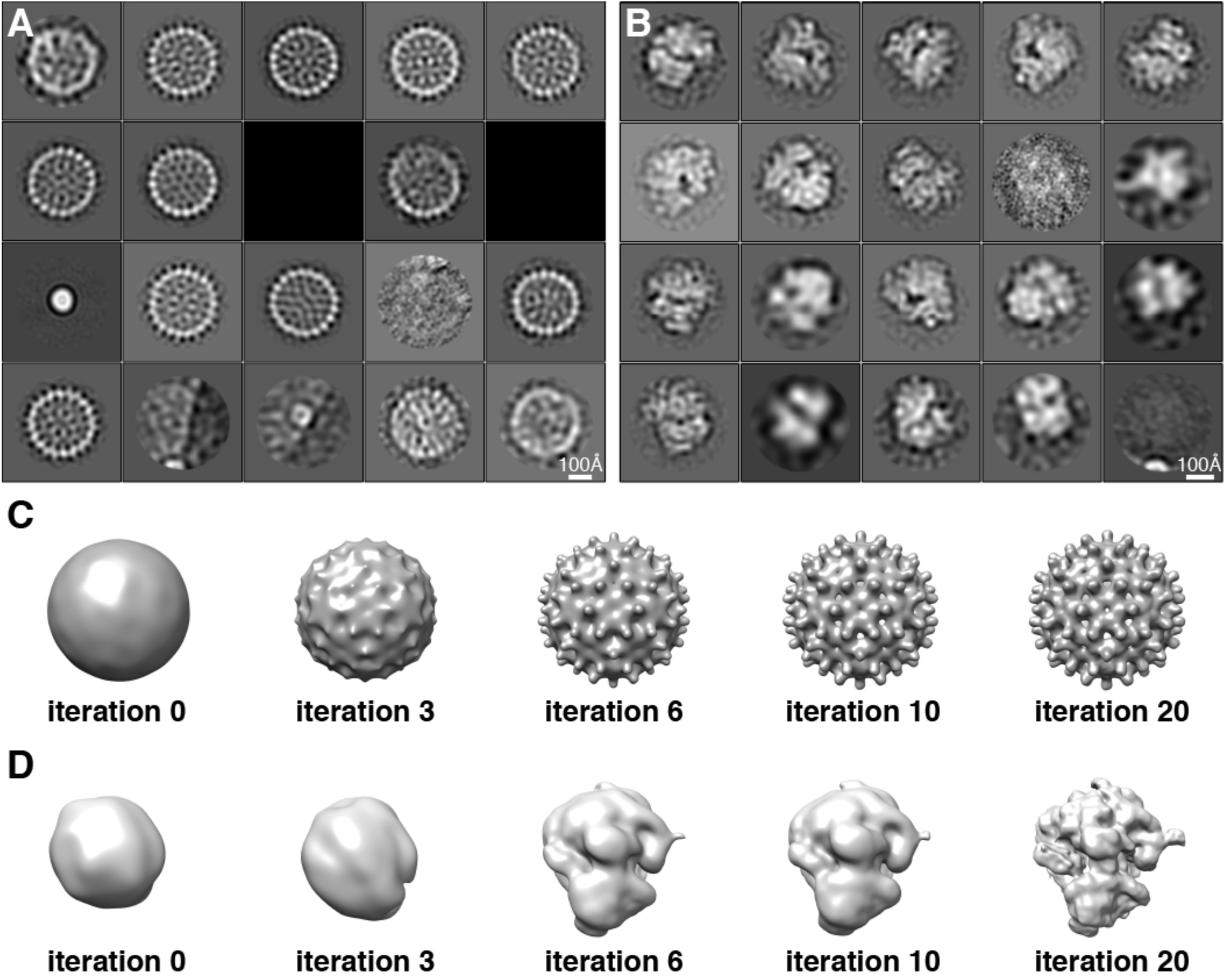
2D Classification and initial model generation. (A) 2D classification of projected sub-tomograms of the HBV capsid particles. Particles were selected from a template matching procedure and this classification helps in removing bad particles, for example ones that correspond to 10 nm gold fiducials. (B) 2D classification of projected sub-tomograms of *S. cerevisiae* 80S ribosomes. These data were picked manually in IMOD. (C) Reference-free refinement of the HBV capsid data set. Sub-tomograms were assigned random Euler angles initially (in iteration 0) and then refinement was commenced. (D) Reference-free refinement of the *S. cerevisiae* 80S ribosome particles, again starting from random orientations.

19| If desired, users may also use the selected 2D classes to write a `.star` file containing only the corresponding sub-tomograms. We again provide a python script on the RELION wiki to facilitate this. Run this script from the project directory –

~~~
python relion_2Dto3D_star.py subtomo_proj2d_sel.star particles_subtomo.star
~~~

This script will take two input `.star` files; the first is the `subtomo_proj2d_sel.star` file containing selected particles from step 18 and the second is the `particles_subtomo.star` file containing all sub-tomograms from step 9. This script will write out the subset of the sub-tomograms selected in step 18 into `output_2Dto3D.star`.

20| Once a subset of particles has been identified using steps 18-19, this subset may be used to get a sub-tomogram averaging structure using the ‘**3D auto-refine**’ job-type. The output rootname on the ‘**I/O**’ tab could for example be `./Refine3D/run1`. Whereas single-particle analysis requires an input 3D reference map, sub-tomogram averaging in either the 3D auto-refine or 3D-classification (see next step) job-types may be performed without an initial reference by stating `None` at the ‘Reference map’ entry on the ‘**Reference**’ tab. In the case an external 3D reference is provided, one also needs to indicate whether it is on the correct absolute greyscale. In general, maps created by RELION from the same data will be on the correct greyscale, whereas maps coming from elsewhere may not be. On the same tab, one also needs to provide the resolution of an initial low-pass filter that will be applied to the input 3D map. To prevent model bias, we typically use relatively harsh filters, e.g. in the range of 40-100 Å. On the ‘**CTF**’ tab, one can turn on CTF correction (provided steps 8-14 were performed). If CTF-correction is performed and a 3D map is provided as initial reference, then one also needs to indicate whether the input map has been CTF-corrected. Maps from RELION may be CTF-corrected (depending on whether CTF correction was performed in previous runs), and also for maps that were generated from an atomic model one needs to indicate they have been CTF-corrected (i.e. to reflect that they don’t suffer from CTF artefacts). On the ‘**Optimisation**’ tab, we typically use similar options as for 2D classification runs, with the exception of the regularisation parameter, which is typically set to 4 for 3D auto-refine and classification (see next step). For the options in the ‘**Auto-sampling**’ tab, the angular and offset sampling rates and ranges will only be used in the first several iterations. After that the auto-sampling algorithm will automatically use finer samplings and smaller ranges until the refinement converges (Scheres, 2012b). The default parameters will be suitable for most projects, perhaps with the exception of particles with icosahedral symmetry, for which initial angular sampling rates of 3.7 degrees, and local angular searches from 0.9 degrees may yield better results. Note that the auto-refinement will divide the data into two random half-sets, each of which will be refined independently, so that in the next step “gold-standard” resolution estimates may be calculated (Henderson et al., 2012; Scheres and Chen, 2012). Executing this job will output multiple files for each iteration in the `./Refine3D` directory. The final `./Refine3D/run1_half[1, 2]_class001_unfil.mrc` files will be used in the post-processing as described in step 24.

21| Once a 3D reconstruction has been obtained using sub-tomogram averaging, the sub-tomograms may be classified using the ‘**3D classification**’ job-type on the GUI to detect different conformational states of the specimen. On the ‘**I/O**’ tab select the original `particles_subtomo.star` file, or the `output_2Dto3D.star` from the 2D classification described in step 19. The output rootname could for example be `Class3D/run1`. Because of computational costs, we often use fewer classes for 3D classification than for 2D classification, with typical values in the range of 3-10. On the ‘**Reference**’ tab, use the final reconstruction from step 20 (`./Refine3D/run1_class001.mrc`) as the reference map. This map is now on the correct absolute greyscale, and a similar initial low-pass filter as in step 20 may be applied. On the ‘**CTF**’ tab indicate that the reference has been CTF-corrected (if this was indeed done in step 20). On the ‘**Sampling**’ tab, the default parameters are again suitable for most projects. Executing this job will result in multiple output files for each iteration in the `./Class3D` directory. The final reconstructions for each class are stored in `.mrc` files called `./Class3D/run1_it025_class0??.mrc`. The `./Class3D/run1_it025_model.star` file contains information about the final classes, such as their relative size and the estimated resolution of each reconstruction.

22| The user again needs to decide which of the classes look good. The ‘**Display**’ button on the GUI may be used to show 2D slices through each of the 3D maps (e.g. select. /`Class3D/run1_it025_class001.mrc` to visualize the first class). Visualization of all classes together in UCSF Chimera (Pettersen et al., 2004) is also useful. Once good classes have been identified, one should select the. /`Class3D/run1_it025_model.star` file from the ‘**Display**’ button on the GUI to display (central slices through) the maps of all classes. A `.star` file with the particles belonging to the good classes can then be saved in the same way as in step 18.

23| Sometimes, performing steps 20-22 multiple times, where selected classes from one run are re-classified in a next run, helps to obtain more homogeneous classes. Thus, each class as identified in steps 21-22 should again be refined to high-resolution using the 3D auto-refinement as described in step 20.

24| After completion of the final 3D auto-refinement for each class, the ‘Post-processing’ job-type is used to obtain resolution estimates that have been corrected for the influence of a solvent mask (Chen et al., 2013), to calculate for the modulation transfer function (MTF) of the detector, and to sharpen the final map. On the ‘**I/O**’ tab, one of the two unfiltered half-maps should be provided as `./Refine3D/run1_half1_class001_unfil.mrc`; the output rootname could be set to `run1_post`. On the ‘**Mask**’ tab, the user may provide parameters for the calculation of an automated solvent mask (automask). The average of the two half-maps will be binarized at the specified initial threshold; extended by the provided number of pixels; and finally a cosine-shaped soft edge with the specified width will be added to the mask. The initial threshold value should be chosen such that the automask does not contain isolated white regions in the solvent area. Often a good estimate for the initial threshold value is the threshold at which a display of the `./Refine3D/run1_class001.mrc` map in UCSF Chimera is noise-free in the region around the particle. Once a suitable automask has been created, it can also be provided as input on the ‘**Mask**’ tab instead of calculating a new automask in subsequent Post-processing runs. On the ‘**Sharpen**’ tab, a MTF curve for the detector may be provided. Curves for some detectors may be downloaded from the RELION wiki. For other detectors, the manufacturer may provide MTF curves. If no curve is available, this entry may also be left empty, in which case MTF-correction will be emulated by additional B-factor sharpening. For maps with resolutions beyond 9-10 Å, automated B-factor sharpening (Rosenthal and Henderson, 2003) may be performed. Alternatively, a user-defined value may be provided. Typically, the option to skip FSC-weighting on the ‘**Filter**’ tab is not used. Execution of this step will generate the final map (`./Refine3D/run1_post.mrc`), the automask (`./Refine3D/run1_post_automask.mrc`) and a `.star` file with the applied B-factor, the resolution estimate and the corrected FSC curve (`./Refine3D/run1_post.star`). In addition, the corrected FSC curve will be written out as a file called `.Refine3D/run1_post_fsc.xml`, which can be directly uploaded to the EMDB.

? TROUBLESHOOTING

25| Because many macromolecular complexes are inherently flexible, even after classification data sets will often still contain some extent of structural heterogeneity. This will lead to local variations in resolution in the refined map. To estimate local resolution variations, the ‘**Local-resolution**’ job-type implements a wrapper to the ResMap program (Kucukelbir et al., 2014). On the ‘**I/O**’ tab, again provide one of the two unfiltered half-maps, and provide the range and step size of the resolutions to be tested by ResMap. One typically does not change the default P-value. ResMap will provide much more reliable resolution estimates if one provides a suitable mask. The automask calculated in the previous step typically performs well. The ‘**Run**!’ button will launch the GUI from the ResMap programs. In most cases one can just hit ‘**Continue**’ inside ResMap, i.e. without adjusting any of its parameters. The result is a file called `./Refine3D/run1_half1_class001_unfil_resmap.mrc`, which can be used inside UCSF Chimera to colour the `./Refine3D/run1_post.mrc` map, using menu options `Tools -> Surface Color -> by volume data value`.

26| When a refined map still shows signs of large amounts of structural heterogeneity, i.e. it contains regions of relatively low local resolution, it may be useful to perform a focused classification on that specific region. In that case, one inputs the `./Refine3D/run1_data.star` file into a new 3D classification run as explained under step 21. In this run, one could use a solvent mask that is only white in the disordered region. Also, one could either skip orientational searches (through the corresponding option on the ‘**Sampling**’ tab), or one could perform relatively fine, but local angular searches around the input orientations. Thereby, one may iterate two or more times through steps 20-25.

## Timing

The time taken for the procedure depends approximately linearly on the number of tomograms, the number of sub-tomograms and (inversely so) on the number of processors used on the cluster. Here, for the HBV capsids, we analyzed a dataset containing 15 tomograms (each occupying ~75 Gb of hard disk space). 1851 capsid particles were extracted in boxes of 240^3^ pixels from these tomograms and data was processed on a computing cluster with 4 hyper-threaded 12-core nodes, each with at least 32 Gb of RAM. CTF volume reconstruction (distributed over 15 cores, 1 per tomogram) took 2 hours. Sub-tomogram extraction and projection of the extracted sub-tomograms took 30 minutes using a single core. 2D classification of the data in 20 classes took ~3 hours on 2 nodes. 3D auto-refinement was concluded in 24 hours, using 4 nodes. Postprocessing was performed on a single core in 10 minutes.

For the 80S ribosomes, 7 tomograms were used and 3120 ribosomes were extracted in boxes of 200^3^ pixels and projected into 2D in 20 minutes using a single core. CTF volume reconstruction took 30 minutes on all nodes; 2D classification into 20 classes took 30 minutes on 2 nodes; 3D classification into 3 classes took 10 hours on 4 nodes; and 3D auto-refinement took 50 hours on 4 nodes.

## Troubleshooting

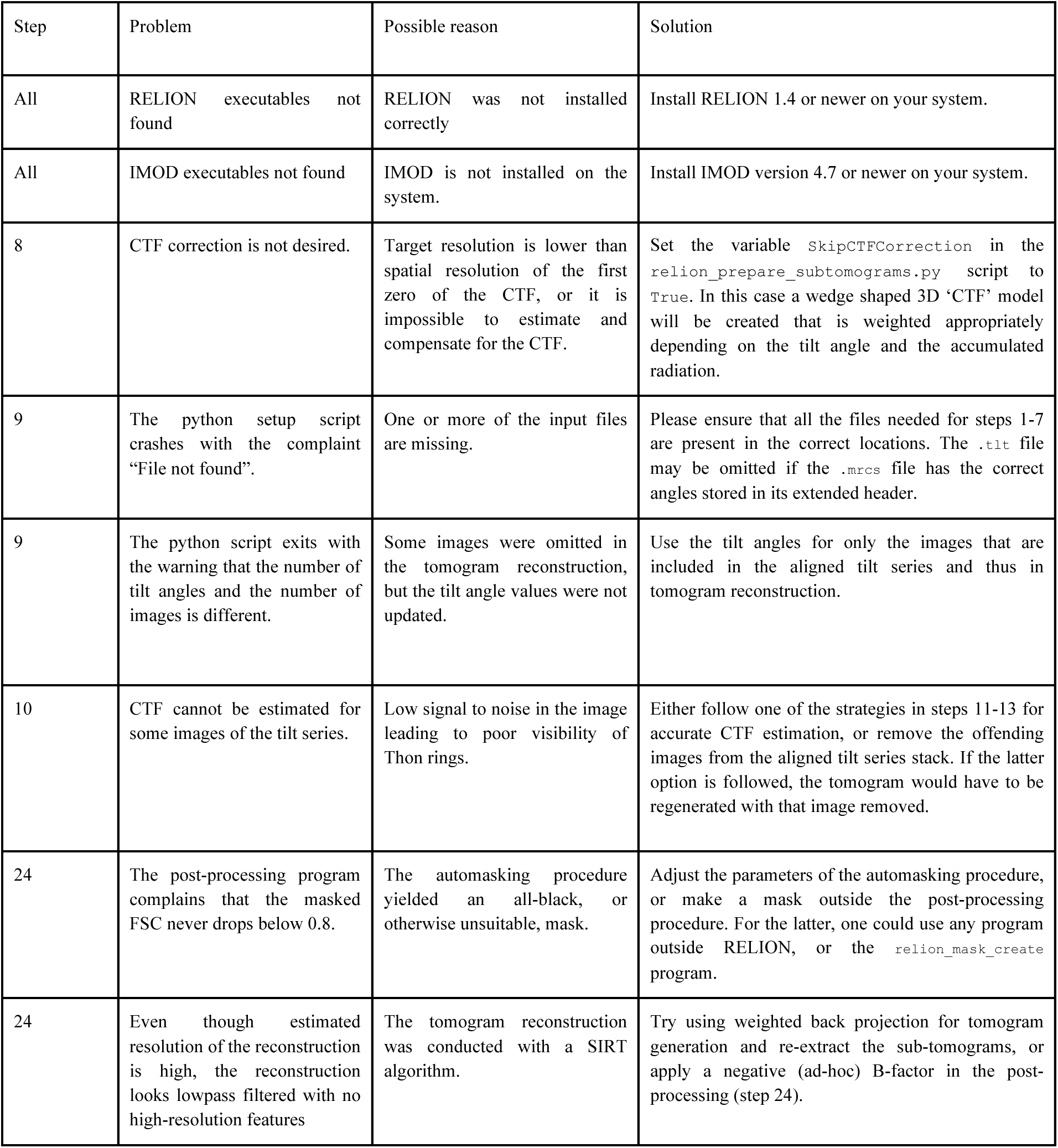

## Anticipated results

We illustrate the results of this protocol for two test data sets. The first set comprised 15 tomograms that were collected on a sample of purified hepatitis B virus (HBV) capsids. For this data set, two extra trial images on either side of the region of interest were used for CTF estimation. After optimizing the input parameters for CTFFIND (defocus search range, resolution search range and box size), CTF estimation from these extra images was found to be adequate even at high tilts (Figure 2D). Thus, the arithmetic mean of the defoci were applied to the tilt series image at every tilt. The coordinate files for the HBV capsids were obtained by automated picking using the template matching routines in the program MolMatch (Förster et al.). In addition to true HBV capsids, this program also picked up 10 nm gold fiducials and other undesirable features. Using both steps 15 and 16, HBV capsid particles were extracted in 240^3^ pixel sub-tomograms, as well as the corresponding 2D projections along the Z-axis. 2D classification, as described in steps 17-19 readily separated HBV capsids from false positives of the automated picking procedure (Figure 4A). 3D auto-refinement (step 20), starting from random orientation assignments for all sub-tomograms, followed by post-processing (step 24), yielded a final map to a resolution of 9.4 Å (Figures 4C and 5A) from 1851 particles where secondary structure elements like α-helices were clearly resolved (Figure 5C).

**Figure 5.**
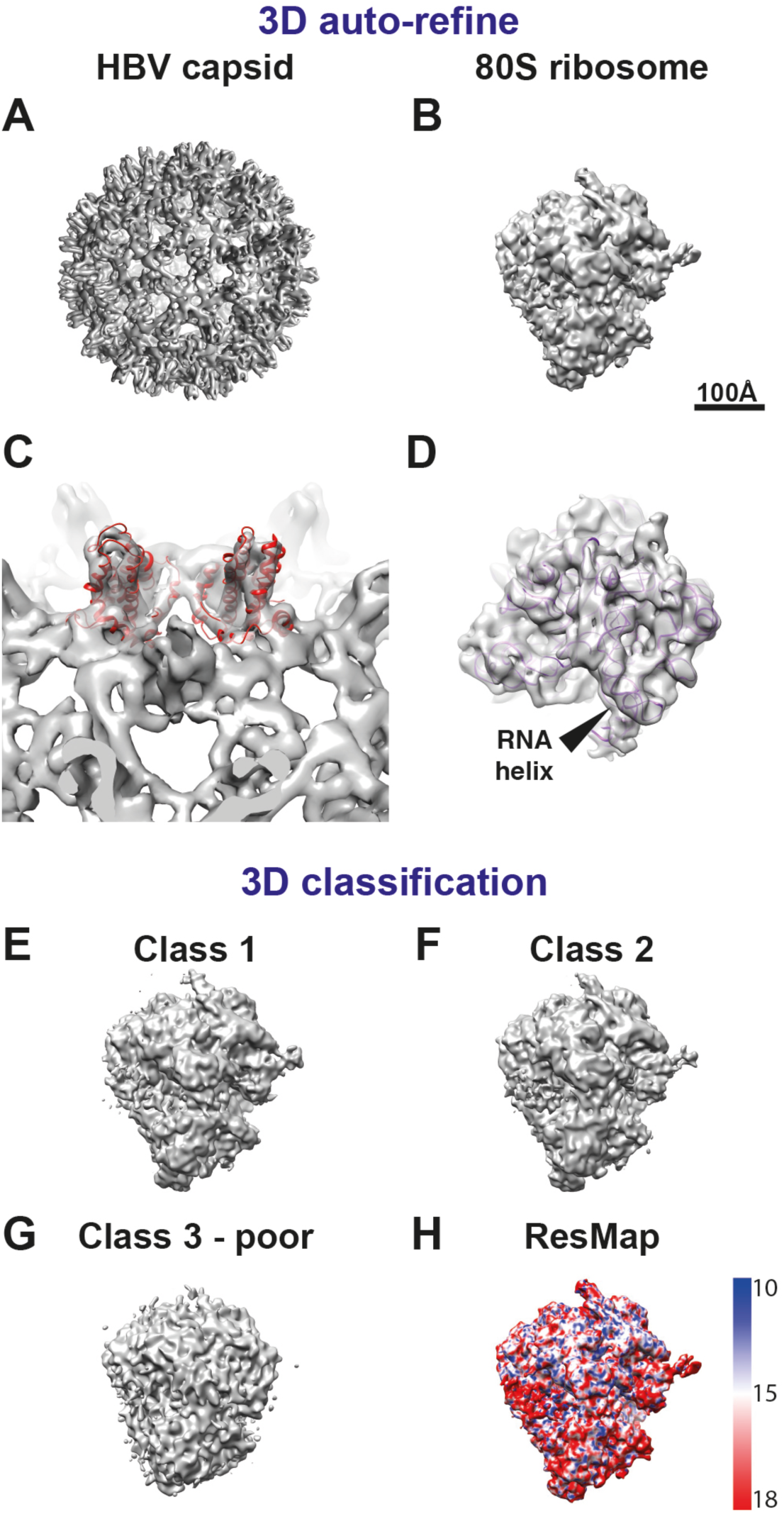
3D auto-refinement and classification using the regularized-likelihood algorithm in RELION. (A) Output of 3D auto-refinement procedure from RELION for the HBV capsid dataset. (B) Output of 3D auto-refinement for the 80S ribosome data set (This map has been deposited at the EMDB under the accession number EMD-3228). (C) Secondary structure features (α-helices) are resolved in the HBV capsid map. Fitted atomic co-ordinates into the sub-tomogram average highlight the positions of the helices. (D) RNA helices resolved in the 80S ribosome map. The atomic co-ordinates have been fitted into this map as rigid bodies for visualization. (E-G) 3D classification of the ribosome data set into three classes reveals a subset of particles (~15% of the data set) in G that show a poor sub-tomogram average. (H) Removing these particles in a second 3D auto-refinement leads to a cleaner map. The result of ResMap is plotted onto the final density showing somewhat lower resolution in the small subunit of the ribosome.

The second test data set of 7 tomograms was collected on a sample of purified 80S ribosomes from *S. cerevisiae.* We have deposited this data set together with the results described here at the EMPIAR data base (Patwardhan et al., 2014) under accession number EMPIAR-10045. The initial tomograms, the corresponding aligned tilt series, as well as the `.tlt`, `.order`, `.trial` and .`coords` files as described in steps 1-7 are all stored in the `./Tomograms` subdirectory of the EMPIAR entry. The results of all other steps are stored in the `./AnticipatedResults` subdirectory of the EMPIAR entry. Thereby, novice users can follow the exact steps described above to replicate, and compare with, the results described here. CTF estimation was conducted using the same procedures as for the HBV capsid data set. However, even after optimizing the inputs of CTFFIND, CTF estimation from high-tilt images was inaccurate in some instances. We believe that this is due to increased specimen thickness due to the deposition of a layer of carbon on the grid during sample preparation. Thus, we used the average of the defoci from the lower tilt images and applied this average to all high tilt images in step 11. Manually picked particles were extracted as sub-tomograms, as well as 2D projections along the Z-axis. 2D classification of the projected sub-tomograms revealed multiple different views of the ribosomes, and some small classes that needed to be discarded (Figure 4B). In this case, 3D auto-refinement of the input 3120 particles, again starting from random orientations, followed by post-processing, led to a 13 Â reconstruction (Figures 4D and 5B). Typical features like grooves of RNA helices were clearly visible in this map (Figure 5D). Using the output of the 3D auto-refinement we also conducted 3D classification (as in step 21) of the entire data set into 3 classes (Figure 5E-G). Although we could not identify different ratcheted states of the ribosomes, we identified a small subset of particles (Class 3, Figure 5G) that gave rise to a poor average. Removing these particles from the data set and subsequent 3D auto-refinement resulted in a somewhat cleaner output map, albeit at the same measured resolution. To assess local resolution variations in this map, ResMap analysis was performed as explained in step 25. The resulting map shows somewhat lower resolution in the small subunit than in the large subunit (Figure 5H).

## Acknowledgements

We thank Xiaochen Bai, Israel Sanchez Fernandez and K. Vinothhumar for help with sample preparation; Jake Grimmett and Toby Darling for assistance with high-performance computing; Shaoxia Chen and Christos Savva for assistance with electron microscopy; and Jan Löwe for helpful discussions. This work was supported by funds from EMBO (ALTF 3-2013 to T.A.M.B.) and the UK Medical Research Council (MC_UP_A025_1013 to S.H.W.S.).

